# There and back again: a multi-omics tale of thyroid co-expression network rewiring

**DOI:** 10.64898/2026.05.21.726849

**Authors:** Marina Pozhidaeva, Hannah Bussmann, Maike Huisinga, Roland Buesen, Jörg Hackermüller, Sebastian Canzler

## Abstract

The integration of multi-omics data offers unprecedented insight into complex biological systems but presents significant analytical challenges. In this study, we propose a best-practice framework for constructing simultaneous weighted gene co-expression networks (WGCNA) from transcriptomics, proteomics, and metabolomics data. Using a rodent model of thyroid toxicity induced by propylthiouracil (PTU), we analyzed thyroid tissues from control, treated, and recovery groups. We demonstrate that concatenating individually processed omics layers at the sample level—without additional scaling—preserves meaningful correlation structures and reflects best practices for biologically interpretable network construction. Co-expression networks were constructed for each group, revealing extensive disruption of molecular interactions under treatment and partial restoration during recovery. We highlight the complementary strengths of two analytical strategies: module preservation analysis identifies disrupted co-regulatory structures, while differential connectivity analysis detects feature-level rewiring events. As a methodological advance, we introduce a permutation-based approach for calculating feature-specific p-values for differential connectivity (DiffK), enabling robust statistical inference. This strategy uncovered over 4,400 significantly rewired features, many of which showed stable expression, underscoring the added value of network-based analyses. Our findings demonstrate the utility of integrated multi-omics WGCNA and differential network analysis in capturing dynamic, system-wide regulatory changes.

## Introduction

In the era of high-throughput technologies, generating large-scale (multi-)omics datasets, such as transcriptomics, proteomics, and metabolomics, has become routine. While such datasets offer unprecedented insights into biological systems, their complexity poses a significant analytical challenges [1]. In a previously performed toxicology study, we utilized multi-omics datasets derived from an oral exposure experiment in rats treated with propylthiouracil (PTU), a compound known to induce hypothyroidism [2]. Our findings demonstrated that multi-layered analyses could offer deeper insights into molecular mechanisms and pathway responses compared to single-layer analyses alone. However, extracting meaningful biological information from such multi-dimensional datasets requires sophisticated methods capable of capturing the intricate interplay among genes, proteins, and metabolites.

Weighted Gene Co-expression Network Analysis (WGCNA) is a powerful systems biology method designed to identify co-regulated expression patterns across biological samples or experimental conditions [3]. By leveraging correlations among molecular features, WGCNA constructs co-expression networks and identifies modules of functionally related molecules, thus providing a holistic perspective beyond single-gene analyses.

Recently, WGCNA-based multi-omics studies have emerged mainly following a sequential approach: each omics layer is analyzed separately, creating independent co-expression networks before identifying and comparing correlated modules across layers [4–6]. There have been individual efforts toward a simultaneous integration, where distinct omics layers are combined and analyzed collectively from the outset. However, such studies remain scarce and lack consensus regarding optimal preprocessing strategies [7, 8]. In general, most existing WGCNA multi-omics studies have integrated two omics layers, with attempts involving three or more layers being notably limited [9].

Preprocessing methods for individual omics layers are typically applied ad hoc, leading to inconsistent outcomes and complicating reproducibility. Several reviews have compared preprocessing strategies, highlighting the variability and emphasizing the absence of universally accepted guidelines [10, 11]. Establishing robust best practices for multi-layer preprocessing and integration is therefore crucial for enhancing the biological relevance and reproducibility of multi-omics analyses.

Network inference enables the investigation of how regulatory networks change in response to external triggers or environmental perturbations. Building on this principle, comparative network analysis emphasizes the exploration of network rewiring across distinct experimental conditions and has been successfully applied in the past [12, 13]. There, changes in connectivity patterns within a network under different biological states are detected, shedding light on condition-specific regulatory relationships [14].

In the present study, we propose a universal and simultaneous multi-omics integration strategy for WGCNA, that can be applied to various omics layers. By integrating transcriptomics, proteomics, and metabolomics data in a unified WGCNA framework, we further seek to uncover cross-omics regulatory relationships that may be missed in isolation. This integrated strategy holds the potential to enhance robustness and biological relevance of network-based analyses. Additionally, we introduce a permutation-based statistical approach to quantify and assess the significance of feature-specific connectivity changes across experimental conditions. By defining robust best practices and exploring network dynamics comprehensively, we provide a powerful analytical framework to elucidate complex biological networks’ structure, dynamics, and function.

## Materials and methods

### Multi-omics data set

The omics datasets used in this study were generated from rodent experiments conducted as part of the CEFIC LRI Project ‘XomeTox’^1^ and have been published in [2]. In the presented work, we focus on thyroid samples and direct thyroid toxicity induced by propylthiouracil (PTU), which was administered via the diet. The experimental design consisted of the control groups, low dose (5 ppm PTU), and high dose (50 ppm PTU) treatment groups. Five animals per group were sampled at three time points: after 2 and 4 weeks of treatment, and after 4 weeks of treatment followed by a 2-week recovery period. The experiment followed a paired-sample design meaning that multiple omics layers were generated from the same tissue. For multi-omics network construction, we used transcriptomics, proteomics, and tissue metabolomics data from the thyroid samples. The transcriptomics data is accessible at NCBI SRA with BioProject identifier PRJNA695243^2^. The proteomics data is available on PRIDE [15] with the ID PXD026835^3^. The metabolomics data is available at Metabolomics Workbench [16] with StudyID ST002023^4^. A range of clinical and pathological parameters were assessed, including body weight, organ weights, and thyroid hormone levels (T3, T4, and TSH). Histopathological evaluation of thyroid tissue focused on follicular hypertrophy and hyperplasia. For a complete description of the experimental procedures, data generation, and assessed endpoints, we refer to [2].

### Data Preprocessing and Comparison of Correlation Patterns

For transcriptomics, we retained genes measured in at least 40 of 45 samples and excluded low-variance genes using a 30th quantile threshold. The variance stabilizing transformation (VST) was applied using the `vst()` function from the `DESeq2` R package [17].

The proteomics data included proteins detected in at least 40 samples, with missing values imputed at 20% of the row minimum. No further transformation was applied as the data was normally distributed and showed no heteroscedasticity.

For the metabolomics data, metabolites detected in at least 40 samples were retained and missing values were imputed using 20% of the row-minimum. To address heteroscedasticity, we applied the VST using the `justvsn()` function from the `vsn` R package [18].

In an alternative approach, we processed all three individual omics layers following the guidelines described in [7]. Since proteomics was not covered, we processed this data set similar to the metabolomics data. Subsequently, all three layers were normalized using *log*_2_(*x* + 1).

To compare the correlation pattern between both approaches, we calculated the biweight midcorrelation of all feature pairs using the `bicor()` function from the `WGCNA` R package [3], offering greater robustness than the Pearson correlation.

### Network construction and comparative network analysis

We constructed multi-omics networks from transcriptomics, proteomics, and tissue metabolomics datasets, each preprocessed according to best-practice recommendations for the respective omics layer. The three layers were concatenated at the sample level and subsequently divided into subsets including control samples, PTU-treated samples (two and four weeks), and recovery samples, comprising 15, 20, and 10 samples, respectively. The minimum recommended sample size for WGCNA is 15; therefore, it is important to interpret the recovery network with caution^5^. Signed co-expression networks were built for each subset with the `blockwiseModules()` function using biweight midcorrelation and identical parameter settings to ensure comparability. The networks were constructed in a single block by tuning the *maxBlockSize* parameter to the number of features. We identified a beta value of 14 using the `pickSoftThreshold()` function to ensure a scale-free topology for all networks. The remaining parameters were set to *minModuleSize* = 100, *mergeCutHeight* = 0.25, and *deepSplit* = 2. By correlating traits with module eigengenes (first principal component of a module), we explored module connections to histopathological and clinical traits, such as hypertrophy, hyperemia, thyroid hormones, organ weights, and PTU concentrations.

The `modulePreservation()` function from `WGCNA` was used to calculate the composite Z-summary and median rank statistics for all six possible pairwise network combinations. The analysis was based on 300 random permutations of feature labels. Weakly preserved modules were defined as 2 *<* Z-summary *<* 10, while non-preserved modules exhibit a Z-summary score below 2 [14].

For the three networks, we performed an overrepresentation analysis on `KEGG` and `Reactome` pathways across all modules and all three layers separately. P-values of the three omics layers were combined using Stouffer’s method [19] and subsequently corrected for a false discovery rate of 0.05 using Benjamini-Hochberg [20].

To assess differential network connectivity, we computed node-wise normalized connectivities for each network using `softConnectivity()` and the maximum network connectivity as denominator, following [21]. Differences in connectivity (*DiffK*) were obtained by subtracting normalized connectivity across three comparisons: Control - Treated, Treated - Recovery, and Control - Recovery. The test statistic was sampled by randomly permuting sample labels 100 000 times. P-values were calculated using the `permp()` function from the `statmod` package [22].

Pathway enrichment was performed utilizing the `multiGSEA` R package [23] for `KEGG` and `Reactome` pathways. Features were ranked based on their `DiffK` value and significance: *DiffK* · (− log_10_(p-value)). The *nPermSimple* parameter was set to one million.

### Network Rewiring analysis

Eleven thyroid-relevant features were chosen among the top differentially connected entities in our multi-omics DiffK analysis and analyzed for connectivity changes under PTU treatment. For each feature, we extracted its 300 strongest neighbors in the Control, Treatment and Recovery networks; the union of these neighbors defined a comprehensive set of nodes. We then assembled an adjacency matrix capturing each feature–neighbor association across the three conditions. These three-dimensional adjacency profiles were clustered by k-means (k = 4) to reveal major rewiring patterns. Each cluster underwent GO over-representation analysis using the R package `clusterProfiler` [24]. Finally, we reconstructed condition-specific subnetworks by extracting feature–neighbor edges from individual clusters. Network visualization was conducted with `Cytoscape` [25].

### Data availability

All data and analysis workflows are publicly available at Zenodo^6^. This repository includes input datasets, R Markdown workflows, over-representation analysis results, the supplementary material file `Pozhidaeva et al Supplementary material.pdf`, and a containerized computational environment providing the exact R version and all package dependencies required to reproduce the analyses.

## Results

### Best practices for multi-omics network construction

To define a suitable preprocessing strategy for a simultaneous multi-omics data integration using WGCNA, we compared preprocessing and integration approaches for transcriptomics, proteomics, and metabolomics data with respect to their effects on variance structure and correlation patterns.

#### Transcriptomics

WGCNA was originally developed to construct weighted gene correlation networks from microarray expression data [3]. However, it can also be applied to RNA-seq data sets, provided they are well normalized and filtered. RNA-seq data typically follow a negative binomial distribution and exhibit heteroscedasticity, where higher expression levels are associated with greater variance. To account for heteroscedasticity, transformations that stabilize the variance across the range of measurements and reduce the influence of extreme values should be applied. This can be achieved by a variance stabilizing transformation (VST or VSN) [17, 26], the `voom` transformation from the `limma` R package [27], or different variations of root and log transformations [28]. To shape RNA-seq data for WGCNA, the authors’ recommendations^7^ are a stringent filtering of genes, e.g., lowly expressed or low-variance genes, followed by a variance-stabilizing transformation using the `vst()` method from the `DESeq2` R package [17]. Accordingly, we filtered the transcriptomics data for sufficient detection and variance and applied `vst()` prior to network construction.

#### Proteomics

With its potential for large-scale quantitative analysis, proteomics data has been successfully utilized in network-based systems biology approaches [4, 29, 30].

Although technical advances in recent years have driven the development of deep proteomics [31], limited detection sensitivity still poses a challenge, allowing the detection of only a fraction of all expressed proteins [32, 33]. Moreover, proteomic datasets often exhibit a substantial proportion of missing values, typically ranging from 20% to 50% in global LC–MS experiments [34]. The prevalence of missing data introduces complexities that frequently and negatively affect downstream analyses, including `WGCNA` [33]. Addressing missing values is therefore critical; however, assigning undetected proteins a concentration value of zero or excluding them from correlation analyses may introduce bias or cause information loss. Accordingly, a range of imputation methods has been proposed, and recent studies have benchmarked different statistical approaches taking also missing value mechanisms into account, including ‘missing completely at random’, ‘missing at random’, and ‘missing not at random’ [35–37]. Imputation methods generally perform best when the proportion of missing values is low [38], making stringent filtering of proteins with extensive missingness an important preprocessing step. In our analysis, we therefore retained proteins detected in at least 40 of 45 samples and imputed remaining missing values using 20% of the row minimum before network construction.

#### Metabolomics

Like other omics layers, metabolomics has proven instrumental in network-based approaches [39, 40]. Notably, metabolomics shares several characteristics with the previously discussed omics layers, including right-skewed distributions, substantial data sparsity, considerable technical noise, and varying sample sizes. Addressing missing values, a common challenge in metabolomics involves leveraging imputation methods discussed in the proteomics domain [41]. Proposed methods include random forests, singular value decomposition, and related approaches [42, 43], alongside simpler procedures such as replacing missing values with a fixed fraction of the minimum observed value [44]. A prerequisite for successful imputation is stringent filtering, often guided by the ‘80% rule’ in metabolomics, which retains only features detected in at least 80% of the samples [45]. Given the diversity of metabolomics platforms, data transformation and normalization should be adapted to the specific properties of the dataset before network calculation. Among the commonly used approaches, variance stabilizing normalization (VSN), log transformation, and probabilistic quotient normalization (PQN) have been identified as suitable methods [46, 47]. VSN, originally adopted from transcriptomics preprocessing, has also been shown to improve quantitative performance in mass spectrometry-based applications, including metabolomics and proteomics [48]. In our analysis, we retained metabolites detected in at least 40 of 45 samples, imputed remaining missing values using 20% of the row minimum, and applied `justvsn()` prior to network construction.

#### Compilation of Multi-Omics Data Set

A key challenge in constructing multi-omics networks lies in integrating heterogeneous data types while preserving biologically meaningful correlation structures. We therefore preprocessed each omics layer individually according to its data-specific requirements and then concatenated the processed layers at the sample level, assuming a paired-sample design where individual omics layers were generated from the same biological sample.

To evaluate whether additional transformations after integration were warranted, we compared this strategy with an alternative preprocessing workflow adapted from [7]. There, transcriptomics and metabolomics data were preprocessed separately using interquartile range filtering and row-minimum imputation, respectively, concatenated at the sample level, and subsequently subjected to log_2_(*x* + 1) transformation and standardization. However, in a three-layer integration scenario, applying these transformations post-concatenation would also affect proteomics data, which exhibit substantially lower heteroscedasticity and do not require additional transformation (Figure 1A). While *log*_2_(*x* + 1) transformation is a common choice for stabilizing variance in metabolomics data, our data indicate that VSN is more effective in mitigating heteroscedasticity in both transcriptomics and metabolomics layers, see Figure 1 B and C.

**Fig 1.**
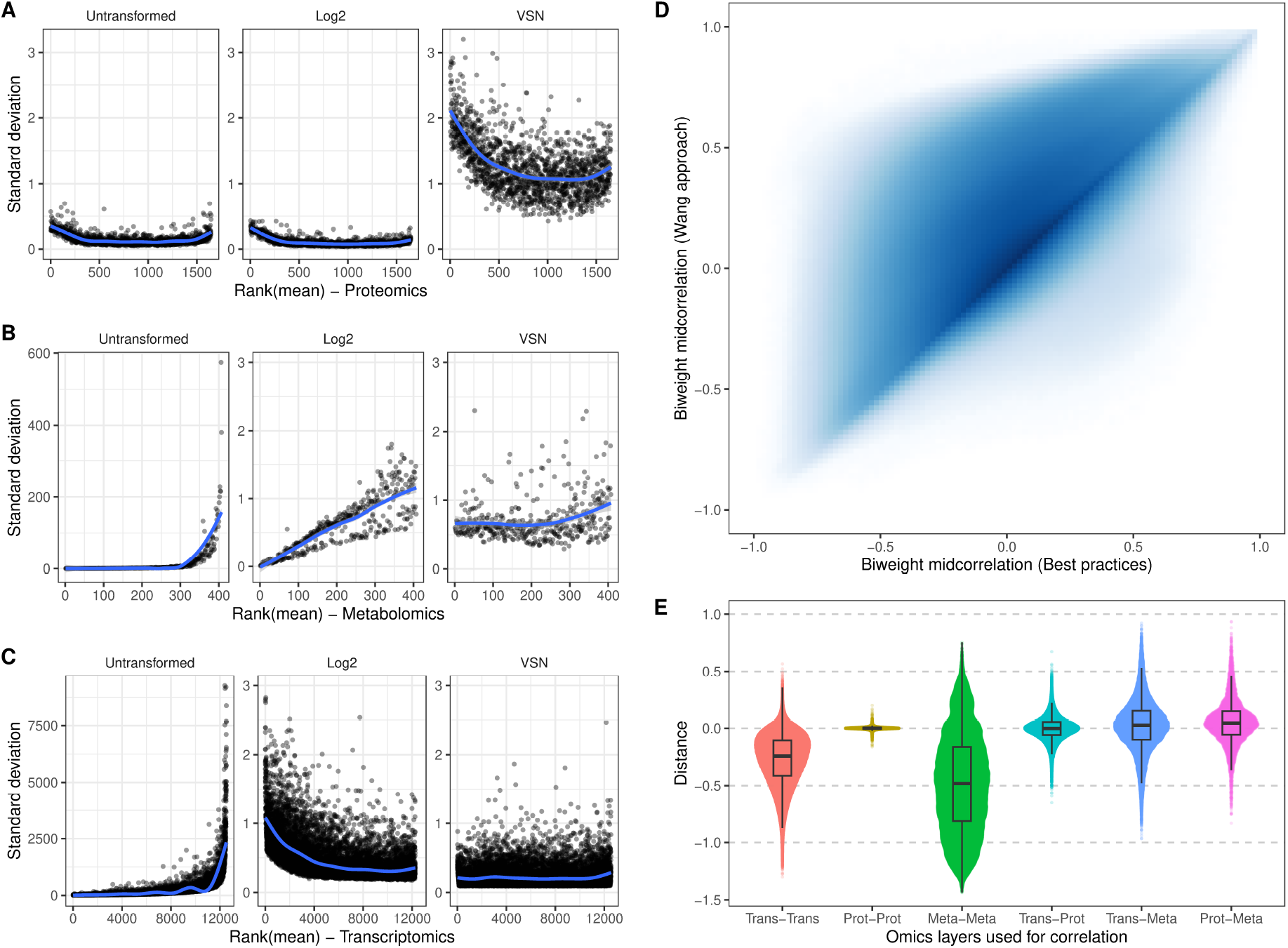
Effects of data normalization and multi-omics layer combination on correlation structure. (A-C) Effects of data normalization to address heteroscedasticity in transcriptomics (A), proteomics (B), and metabolomics data (C). (D) Comparison of the correlation structure following the best practices on individual omics layers and the approach that was introduced by [7]. Biweight midcorrelation was used to calculate all pairwise correlation coefficients for the 14,604 features in the multi-omics data set. Details on the construction of the multi-omics data set are given in the text. (E) Distances between biweight midcorrelation coefficients of both approaches (bicor(Best)-bicor(Wang)) summarized across omics layer pairs.

We next compared the resulting correlation structures between both workflows. This revealed substantial differences in correlation distributions both within and across omics layers (Figure 1D,E). These differences were particularly pronounced in transcriptomics and metabolomics correlations, including gene-gene, metabolite-metabolite, and gene-metabolite correlations and were primarily attributable to the different transformation strategies applied. In contrast, protein-protein correlations were highly similar between both workflows, indicating that the additional log transformation had only minor effects on the proteomics layer.

Because centering and scaling are linear transformations and therefore do not alter Pearson or biweight midcorrelation coefficients [49], additional standardization after concatenation did not provide a benefit for network construction; the mathematical justification is given in Supplementary Section S1.

Based on these results, we advocate for sample-wise concatenation of individually preprocessed omics layers without further scaling for all downstream multi-omics network analyses. Our approach offers several advantages:

1. **Adherence to best practices:** By applying variance stabilizing normalization to transcriptomics and metabolomics data, we more effectively address heteroscedasticity and retain meaningful correlations.
2. **Minimization of transformations:** Reducing unnecessary data transformations mitigates the risk of introducing biases or artificial correlation distortions.
3. **Avoidance of standardization:** Linear transformation and standardization have no impact on the underlying correlation structure.

In summary, our proposed method prioritizes biological fidelity by leveraging best practices for individual omics layers while maintaining a straightforward and interpretable integration strategy.

### Characterization of Multi-Omics Networks and Trait Associations

Following the best practices defined above, each single-omics layer was preprocessed individually, and the three layers were subsequently concatenated at the sample level. The combined dataset was then divided into control, PTU-treated, and recovery groups (two weeks post-treatment), comprising 15, 20, and 10 individual samples, respectively. The networks contained 14,604 features, including 12,550 genes, 1,647 proteins, and 407 metabolites. To ensure dataset integrity, hierarchical clustering confirmed the absence of outliers (Supplementary Figure S1).

Figure 2 shows the scale-free topology model fit (A) and mean connectivity (B), which guided the selection of a soft-thresholding power of 14 for all three networks. This ensured comparable network construction while maintaining scale-free topology (signed *R*^2^ ≥ 0.8) and sufficiently high mean connectivity.

**Fig 2.**
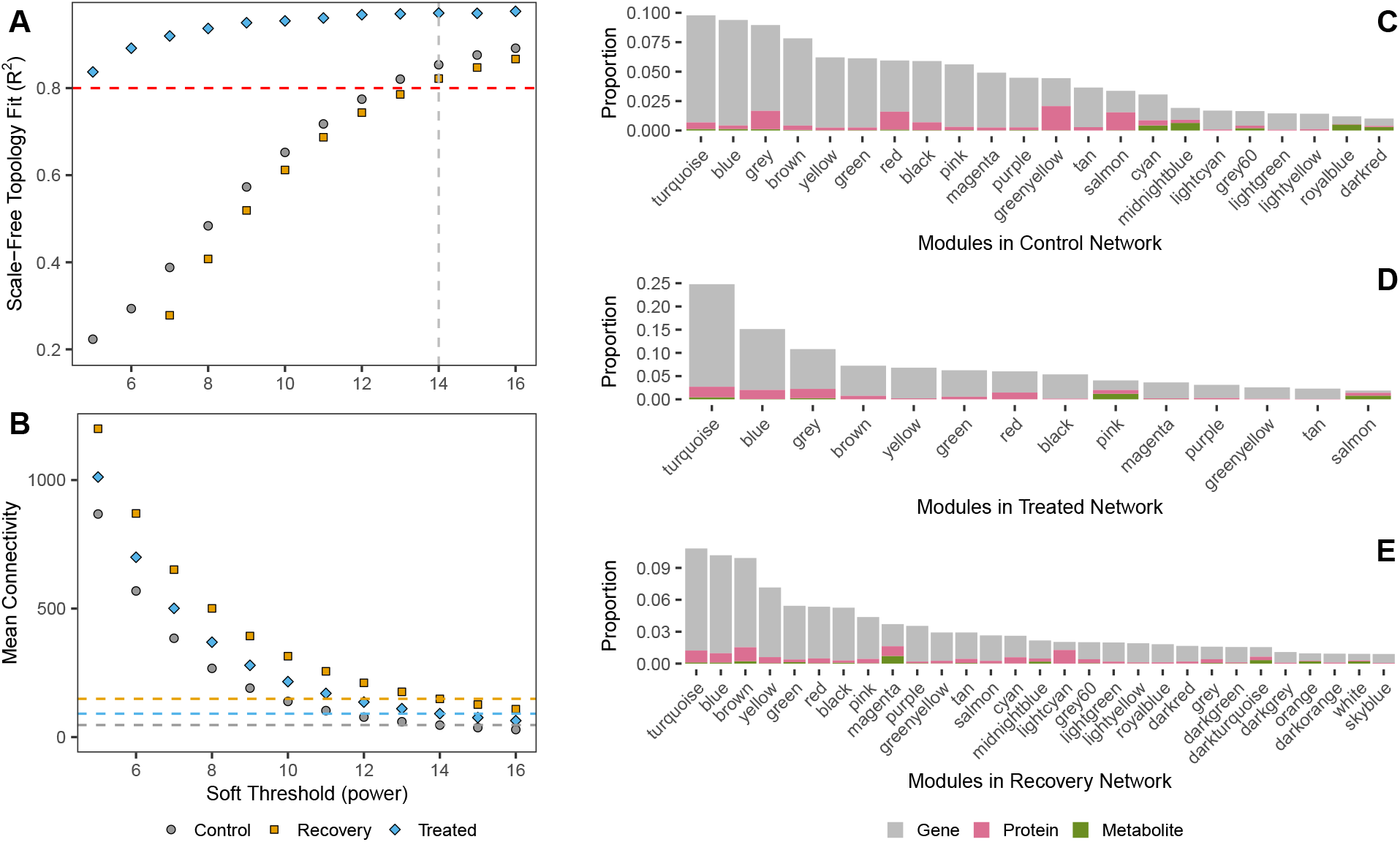
Construction of Weighted Co-Expression Network: Soft Threshold Selection and Molecular Feature Distribution. (A) Scale-Free Topology Model Fit: Evaluation of the network’s adherence to a scale-free topology for various soft-thresholding powers. (B) Mean Connectivity: Average connectivity of nodes within the network across different soft-thresholding powers. (C-E) Molecular Feature Distribution: Proportion of different molecular types (genes, proteins, metabolites) relative to the dataset in the control, treated, and recovery networks.

The WGCNA workflow identified 21 modules in the control network, 13 in the treated network, and 28 in the recovery network. Unassigned features numbered 1,308, 1,579, and 232 in the control, treated, and recovery networks, respectively. Module sizes ranged 148–1,428 features in the control network, 277–3,620 in the treated network, and 132–1,581 in the recovery network.

The hierarchical clustering dendrograms of the features and their module assignments are illustrated in Supplementary Figure S2.

We assessed the distribution of the multi-omics features — genes, proteins, and metabolites — within each module and network. All modules were found to contain features of all three types. Generally, the proportion of each feature type varies slightly between modules but aligns with the overall dataset ratio, as depicted in Figure 2C-E.

Heatmaps illustrating the correlations between module eigengenes and external traits for the three networks are presented in Figure 3, limited to modules that exhibited at least one statistically significant association. In the control network, only 10 of 21 modules displayed significant correlations with at least one trait. The red module exhibited a negative correlation with the time parameter. The lightgreen and greenyellow modules showed positive correlations with thyroid-stimulating hormone (TSH) levels, while the salmon module demonstrated a negative correlation with TSH. In the treated network, the two largest modules, turquoise and blue, exhibited high correlations to thyroid hormone levels, relative thyroid weight, and PTU concentrations. The red module correlated with treatment duration and showed a weaker association with T3 levels. In the recovery network, the red module negatively correlated with PTU concentration and relative thyroid weight, while the yellow module showed positive correlations with PTU concentration. Full heatmaps encompassing all modules are provided in the Supplementary Figure S3. Please note that module colors are assigned independently within each network; therefore, identical colors across networks do not imply similarity or correspondence between modules.

**Fig 3.**
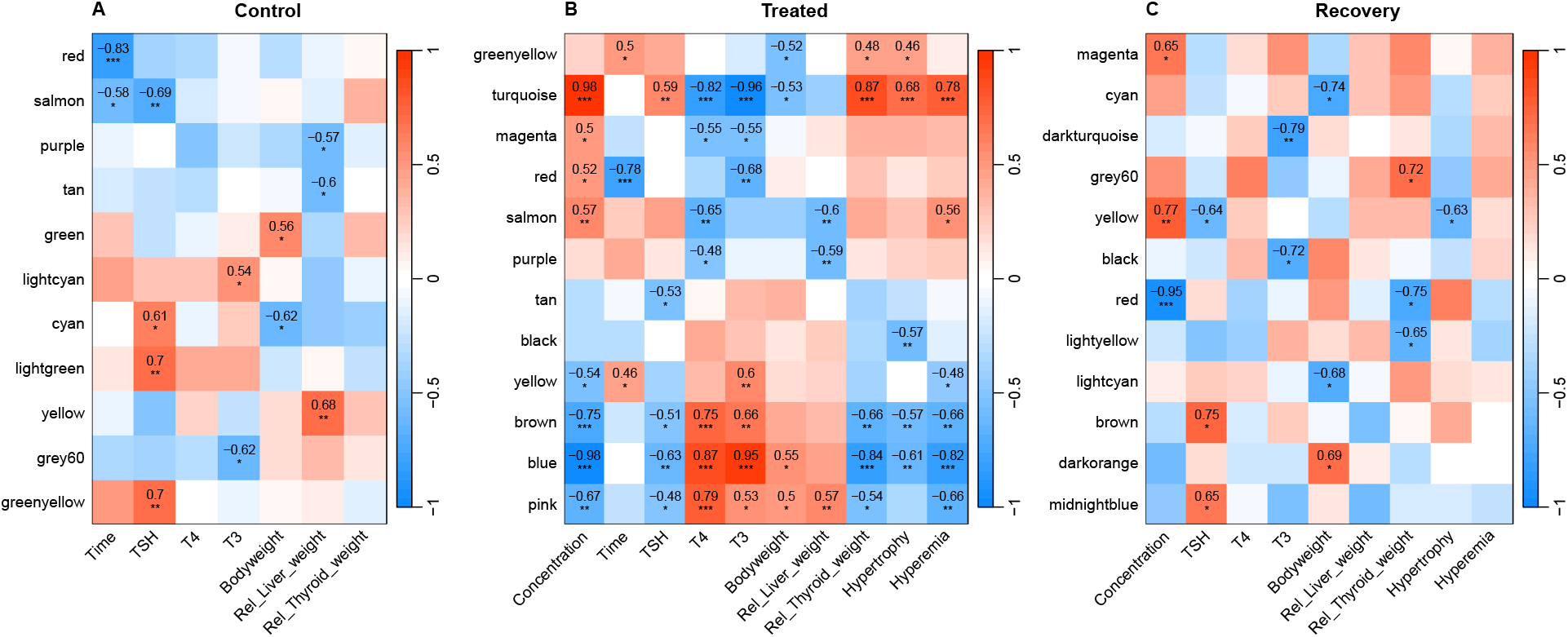
Association analysis of gene co-expression network modules with experimental, clinical, and histopathological traits. The figure displays representative module–trait heatmaps, limited to modules that exhibited at least one statistically significant correlation with external traits across the three networks. Histopathological traits are excluded from the control network matrix due to the absence of hypertrophy and hyperemia, and concentration data is not included for the control group. The time variable is not included in the recovery matrix as all animals were sacrificed after 6 weeks. Module colors are assigned independently within each network; therefore, identical colors across networks do not imply similarity or correspondence between modules. Asterisks indicate statistical significance: * for *p <* 0.05, ** for *p <* 0.01, and *** for *p <* 0.001. Full heatmaps encompassing all modules are provided in Supplementary Figure S3.

### Module preservation analysis

Module preservation analysis is a powerful approach for evaluating module stability across different network conditions, highlighting weakly preserved or disrupted modules. The composite Z-summary score provides an absolute measure for classifying the module preservation status (see Figure 4). The median rank offers relative insights that are less influenced by module size (see Supplementary Figure S5; [14]).

**Fig 4.**
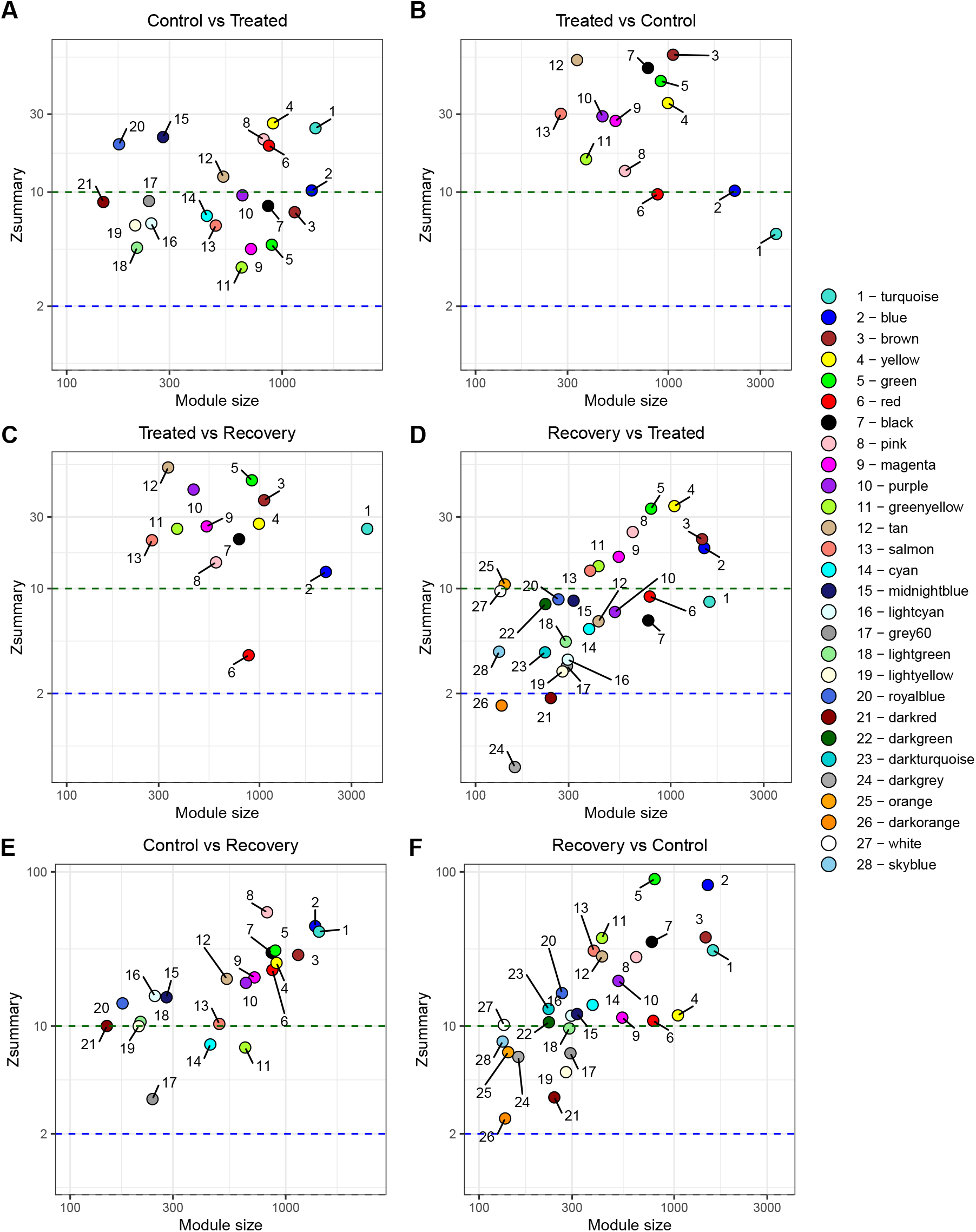
Module Preservation Analysis. (A–F) Z-summary statistics for module preservation across six comparisons. Points represent modules, color-coded by their names, with numeric labels. Horizontal dashed lines indicate Z-summary thresholds of 0 (black), 2 (blue), and 10 (dark green), with modules above the dark green line considered strongly preserved.

Of the 21 modules in the control network, eight were preserved in the treated network (Z-summary ≥ 10), while 13 showed moderate preservation (2 ≤ Z-summary *<* 10) (Figure 4A). The greenyellow, magenta, and lightgreen modules had the lowest Z-summary scores (3.5, 4.5, and 4.6, respectively), suggesting these co-expression patterns were particularly affected by PTU. Notably, those modules also showed a substantial amount of features (∼ 13%) that lost their assignment in the treated network, meaning that they were not co-expressed with other features anymore. Cross-module assignments are shown in the heatmaps in Supplementary Figure S4.

Among the 13 modules in the treated network, 11 were preserved in the control network (Figure 4B), suggesting that most treatment-induced co-expression relationships reflected existing biological structures rather than novel, transient states. The large turquoise module, associated with PTU concentrations and thyroid hormones, and the red module showed moderate preservation with Z-summary scores of 5.5 and 9.7, respectively. The red module, however, had the lowest preservation score (3.6) when analysed in the recovery network. Although 98% of its features remained assigned to modules in recovery networks, they were redistributed among other modules without significant overlap, suggesting that while its individual components persisted, their co-expression patterns changed (Supplementary Figure S4B). The remaining 12 modules showed strong preservation evidence, reinforcing the notion that despite treatment, the majority of network structures remained resilient.

Of the 28 modules in the recovery network, three (darkgrey, darkorange, and darkred) lacked preservation in the treated network, while 16 were only moderately preserved. The least preserved among the moderately stable modules were lightyellow, grey60, lightcyan, darkturquoise, skyblue, and lightgreen, with many of these also displaying weak preservation in the control network. The orange module was preserved in the treated network but not in control, indicating a treatment-specific reorganization that did not persist into recovery. Conversely, most recovery modules were preserved in the control network, mirroring the strong preservation of control modules in recovery. This suggests a broad return to baseline co-expression patterns post-treatment. Some modules, however, showed lower preservation indicating an incomplete recovery for certain processes, e.g., grey60, greenyellow, and cyan with Z-scores of 3.4, 7.3, and 7.6, respectively.

To biologically interpret module preservation and assess whether preservation status correlates with functional significance, we performed overrepresentation analysis (ORA) using `KEGG` and `Reactome` pathways. Enrichments were calculated for each omics layer separately, subsequently combined using Stouffer’s method [19] and adjusted for multiple testing using Benjamini-Hochberg [20]. The false discovery rate threshold for the combined p-values was set to 0.05. Summing across all modules within each network, we identified 79 unique `KEGG` pathways and 282 unique `Reactome` pathways in the control network, 91 `KEGG` and 504 `Reactome` pathways in the treated network, and 83 `KEGG` and 403 `Reactome` pathways in the recovery network. No pathway was enriched across all three omics layers in any network. In the control network, five pathways were significantly enriched in two layers; in the treated network, this number increased to eight; and in the recovery network, no pathway showed enrichment in multiple layers. The complete enrichment results are available on Zenodo^8^.

To pinpoint biological functions disrupted by PTU, we examined the modules with weak or no preservation across networks. In the control network, the cyan module was enriched for energy and lipid metabolism pathways and lost coherence under treatment and recovery. Likewise, the turquoise module of the treatment network, which is poorly preserved in control, harbors a similar set of metabolic processes. Together, these observations point to a condition-specific rewiring of energy homeostasis and lipid handling under PTU treatment, consistent with altered mitochondrial function and fatty-acid metabolism as a direct consequence of reduced thyroid hormone signaling [50, 51]. The red module in the treatment network, being correlated with treatment duration, PTU concentration and T3 levels, was weakly preserved in both control and recovery. There, we identified pathways linked to oxidative stress response, genomic stability, DNA damage repair, and mitotic cell division control. This suggests that extended and increased PTU exposure not only remodels metabolic changes but also induces cellular stress and impacts genomic stability and cell cycle regulation. An overview of enriched pathways in these weakly preserved modules is given in Table 1.

**Table 1.** Summary of pathway enrichment in weakly preserved modules across the control, treated, and recovery networks. Each row corresponds to a module associated with biologically relevant processes, annotated with linked phenotypic traits and reference networks.

| Test Network | Module | Reference Network | Traits | Pathway Categories |
| --- | --- | --- | --- | --- |
| Control | Greenyellow | Treated, Recovery | TSH | Sphingolipid metabolism (2); Glycosaminoglycan catabolism (2) |
| Control | Lightgreen | Treated | TSH | Sphingolipid biosynthesis (1) |
| Control | Lightcyan | Treated | T3 | N-linked glycoprotein biosynthesis and quality control (3) |
| Control | Cyan | Treated, Recovery | TSH, BW | Mitochondrial energy metabolism and biogenesis (16); Lipid metabolism and transport (26) |
| Treated | Turquoise | Control | PTU, T3, T4, TSH, BW, RelThW, HiPa | Mitochondrial energy metabolism and biogenesis (16); Lipid metabolism and transport (9) |
| Treated | Red | Control, Recovery | Time, T3 | Genomic stability and DNA damage response (32); Cell division and mitotic regulation (19); Oxidative stress response (6); p53 activity (6) |
| Recovery | Darkturquoise | Treated | T3 | Lipid metabolism and transport (26) |

**Table notes.** Trait codes: Time – treatment duration; TSH – thyroid-stimulating hormone; T3 – triiodothyronine; T4 – thyroxine; PTU – PTU concentration; BW – bodyweight; RelThW – relative thyroid weight; HiPa – histopathological features (hypertrophy, hyperemia). Pathway categories were manually curated and grouped based on biological function, not derived from automated clustering. Numbers in parentheses indicate the number of significantly enriched terms per category. Only modules with weak or no preservation across networks are included. Full enrichment data are available at Zenodo: ORA\_results\_multiomics\_WGCNA.zip.

### Differential connectivity analysis

We employed differential connectivity analysis [21] to quantify changes in feature-level network connectivity across the three multi-omics networks derived from control, treated, and recovery samples. This approach captures differences in co-expression topology and helps identify biologically meaningful network rewiring events. To quantify changes in feature connectivity, we computed the whole-network connectivity for each feature within each network, normalized by the maximum connectivity. The differential connectivity (*DiffK*) was then defined as the difference in normalized connectivity between network pairs. A key methodological advance of our approach is the implementation of a permutation-based significance test for each feature’s DiffK value. By randomly permuting sample labels and re-computing *DiffK* across iterations, we derived feature-specific p-values that allow statistical assessment of differential connectivity at the individual feature level, something that was not previously implemented. This innovation enables robust identification of significantly differentially connected features (DCFs), extending *DiffK* from a descriptive to an inferential framework.

Using our permutation test, we identified a total of 4,482 DCFs between the control and treated networks, including 3,798 genes, 616 proteins, and 68 metabolites. Between treated and recovery networks, 497 DCFs were identified, containing 489 genes and 8 proteins (Figure 5A). However, when comparing the control and recovery networks, we found no significantly differentially connected features. Volcano plots in Figure 5B, highlight different DCFs associated with thyroid-related biological processes.

**Fig 5.**
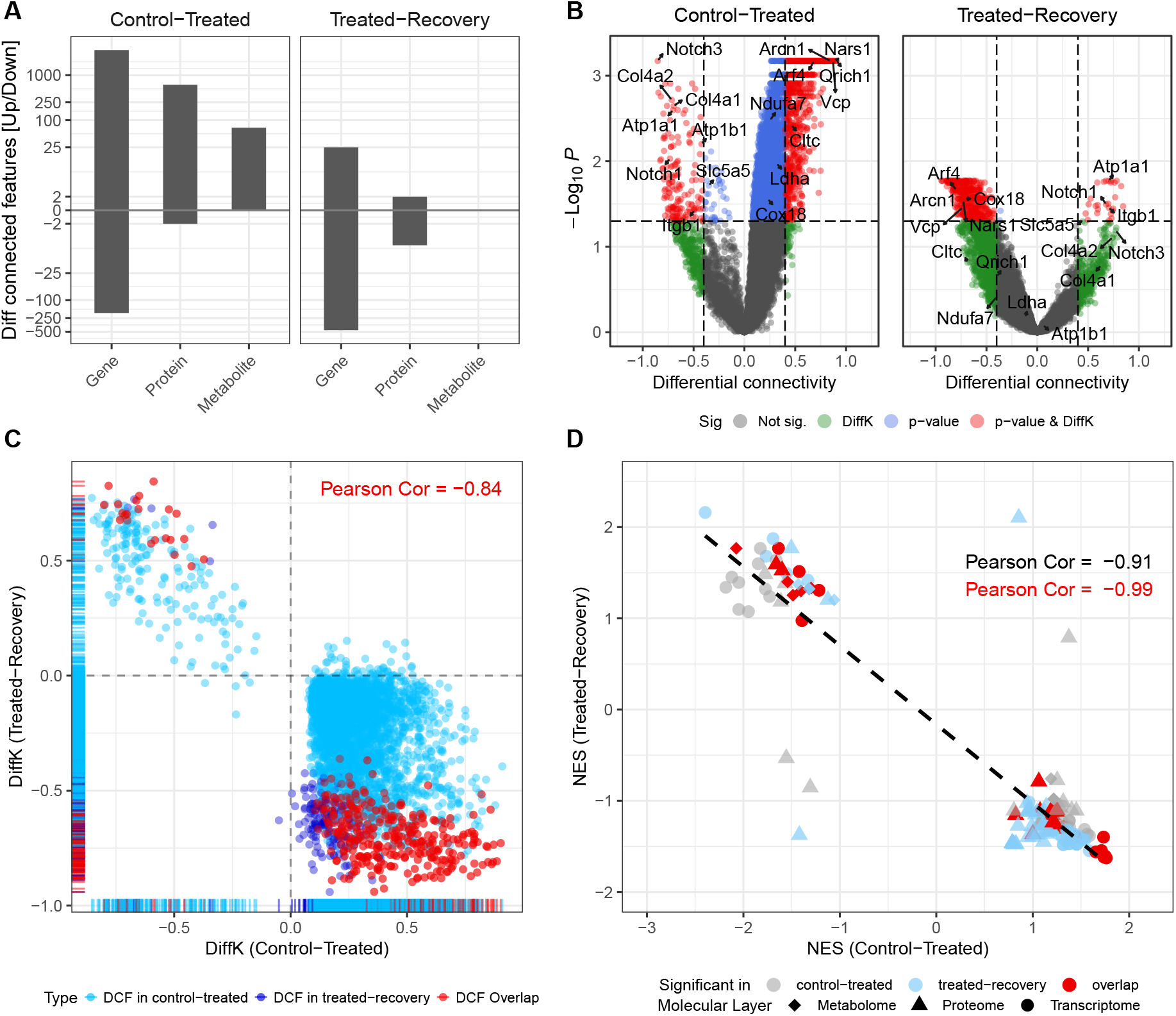
Differential Connectivity Analysis. (A) Bar plot summarizing the number and composition of different molecule types across significantly differentially connected features (DCFs) (FDR ¡ 0.05). The comparison control-recovery is not shown since no DCFs have been detected. (B) Volcano plot highlighting significant features with strong differential connectivity and annotations for those linked to thyroid processes potentially affected by PTU treatment. (C) Scatter plot illustrating the correlation of DCFs between control-treated and treated-recovery conditions. Each point represents a feature, with DiffK values from control *vs*. treated plotted on the x-axis and DiffK values from treated *vs*. recovery on the y-axis. Pearson correlation coefficients indicate the extent of connectivity restoration or further disruption under recovery conditions. (D) Scatter plot comparing normalized enrichment scores (NES) of pathways between control–treated and treated–recovery conditions. Each point represents a pathway enriched in at least one comparison. Shapes indicate the omics layer (transcriptome, proteome, or metabolome) from which the plotted NES is derived. Colors indicate the condition(s) in which the pathway was found to be significantly enriched, based on p-value combination using multiGSEA across omics layers.

Interestingly, the vast majority of DCFs (4,288 of 4,482) exhibited decreased connectivity in the treated network compared to control. In contrast, most DCFs (470 of 497) gained connectivity during recovery compared to treatment, suggesting that PTU treatment disrupts the co-expression structures of a large portion of the biological system, with a partial restoration of those structures upon recovery. This observation is supported by a strong negative correlation (Pearson *r* = −0.84) between *DiffK* values in the control-treated and treated-recovery comparisons (Figure 5C).

By leveraging feature-level p-values from our permutation tests, we next performed a pathway-centric rewiring analysis using multiGSEA. We ranked each feature by a composite score that combines effect size and significance *DiffK* × − log_10_(p-value) and calculated the enrichment for `KEGG` and `Reactome` pathways. This integrative strategy enables identification of differentially connected pathways on a global scale across different omics layers.

We identified 31 significantly enriched pathways in the control-treated (C-T) comparison and 50 in the treated-recovery (T-R) comparison (Table 2). Most enrichments were driven by transcriptomic features. However, we also observed three and five pathways that were individually enriched in the proteomics and metabolomics layer, respectively. The complete lists of enriched pathways can be found in the Supplementary Table S1 and S2.

**Table 2.** Significantly enriched pathways from Reactome or KEGG (marked with ∗) based on differential connectivity results. Adjusted p-values are derived from multi-omics layer combination using `multiGSEA`.

| Pathway | Adj. p (C-T) | Adj. p (T-R) | Group |
| --- | --- | --- | --- |
| ECM-receptor interaction* | 6.170e-04 | 5.802e-01 | C-T |
| The citric acid (TCA) cycle and respiratory electron transport | 7.232e-01 | 1.180e-03 | T-R |
| Collagen degradation | 3.908e-02 | 4.391e-01 | C-T |
| Signaling by NOTCH | 3.472e-02 | 9.145e-01 | C-T |
| ER to Golgi Anterograde Transport | 5.957e-06 | 9.807e-05 | C-T/R |
| Membrane Trafficking | 3.935e-06 | 1.019e-01 | C-T |
| Assembly of collagen fibrils and other multimeric structures | 7.758e-04 | 2.409e-01 | C-T |
| COPII-mediated vesicle transport | 6.082e-03 | 3.341e-02 | C-T/R |
| Cellular responses to stress | 3.103e-03 | 2.516e-01 | C-T |
| Asparagine N-linked glycosylation | 5.654e-07 | 1.042e-05 | C-T/R |
| Vesicle-mediated transport | 1.462e-05 | 8.076e-02 | C-T |
| PTEN Regulation | 3.313e-01 | 3.407e-02 | T-R |
| COPI-mediated anterograde transport | 3.254e-03 | 1.322e-03 | C-T/R |
| COPI-dependent Golgi-to-ER retrograde traffic | 8.165e-03 | 8.076e-02 | C-T |
| Intra-Golgi and retrograde Golgi-to-ER traffic | 1.803e-05 | 8.076e-02 | C-T |
| Golgi-to-ER retrograde transport | 2.500e-04 | 2.421e-01 | C-T |
| Cargo recognition for clathrin-mediated endocytosis | 4.989e-02 | 9.470e-01 | C-T |
| Cellular responses to stimuli | 3.103e-03 | 2.516e-01 | C-T |
| Transport to the Golgi and subsequent modification | 3.935e-06 | 9.807e-05 | C-T/R |

**Table notes.** Pathways with an asterisk (\*) originate from KEGG; others are from Reactome. C-T: Control *vs.* Treated, T-R: Treated *vs.* Recovery, C-T/R: significant in both comparisons. All p-values are adjusted for multiple testing.

Among the top differentially rewired processes, ER to Golgi Anterograde Transport showed extremely strong rewiring in C-T and remained significant in T-R, highlighting persistent remodeling of secretory pathway flux. Likewise, Asparagine N-linked glycosylation and the broader category Post-translational protein modification were among the most dynamically rewired, suggesting that PTU broadly perturbs protein maturation and folding.

In general, pathway-level rewiring mirrored the behavior observed for individual features. Normalized Enrichment Scores (NES) showed a strong inverse correlation between the C–T and T–R contrasts (Figure 5D), indicating that pathways gaining connectivity under treatment tended to lose connectivity during recovery, and *vice versa*. This reciprocal pattern supports the largely reversible nature of PTU-induced network alterations and demonstrates the utility of *DiffK* -based enrichment for identifying dynamically rewired biological processes.

### PTU-Induced Rewiring of Local Feature Networks

To assess how PTU reshapes the local connectivity around major differentially connected features, we constructed feature-centered subnetworks by extracting each feature’s 300 strongest neighbors in control, treatment, and recovery conditions. We focused on features that exhibited significant connectivity gains or losses under PTU treatment (see Figure 6 for the selected hubs). Across conditions, these local feature-specific networks shared only a few edges in common, reflecting the extensive rewiring induced by PTU and the subsequent recovery.

**Fig 6.**
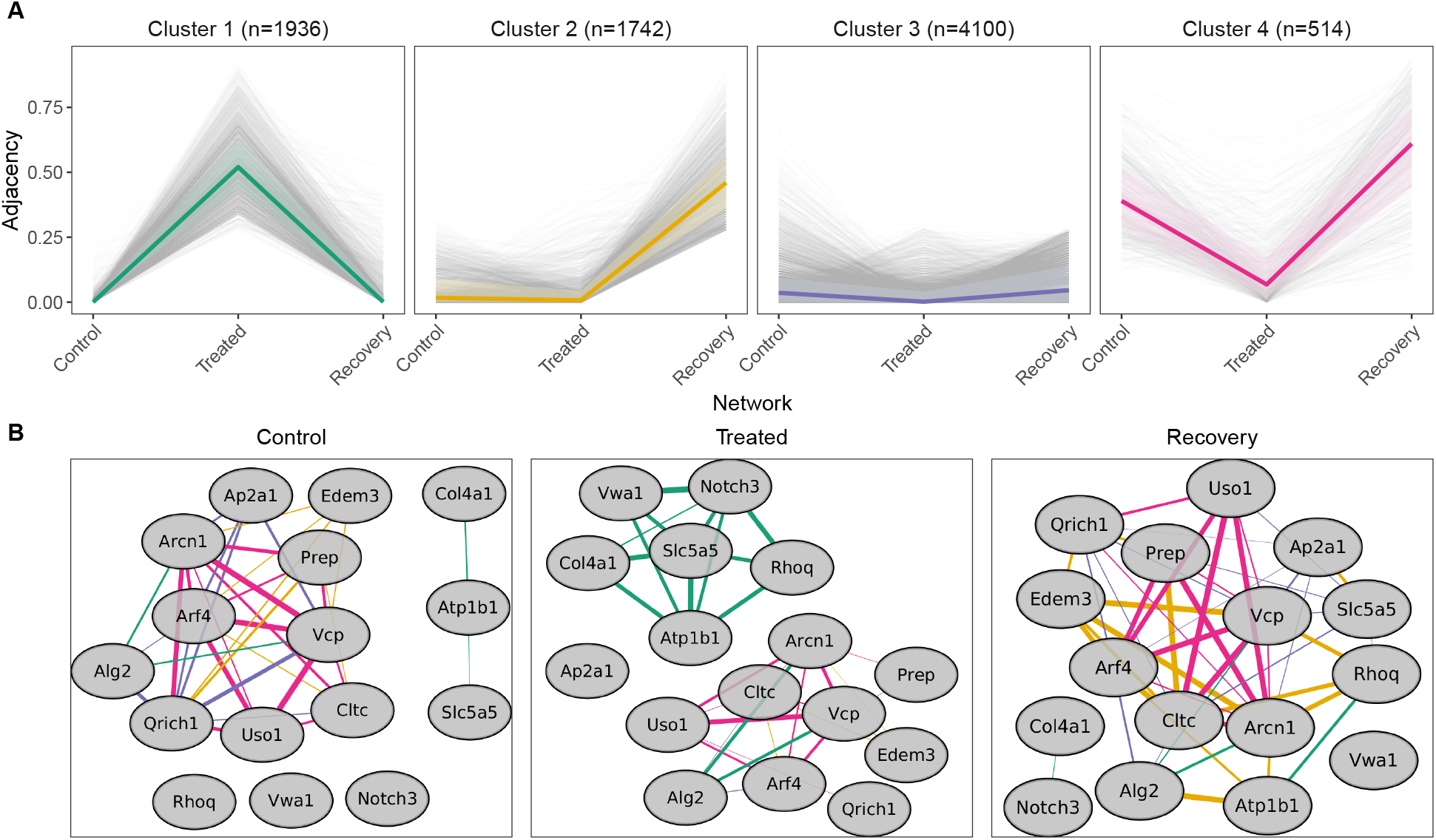
Condition-specific local networks around thyroid-relevant features. (A) Adjacency profiles across Control, Treated, and Recovery conditions, grouped by *k*-means clustering of pairwise connectivity profiles. Grey lines represent individual observations, colored lines indicate median adjacency profiles per cluster, and colored ribbons show interquartile ranges (25th–75th percentile). Each facet represents one cluster with the size indicated in parentheses. (B) Subnetworks were rendered in Cytoscape for each condition using edges derived from selected clusters. Nodes represent thyroid-relevant features, and edge colors correspond to cluster assignments shown in panel A. Edges with adjacency exceeding 0.03 were retained; thickness reflects adjacency.

Clustering the three-dimensional adjacency profiles (k = 4) uncovered four characteristic rewiring trajectories (Figure 6A). Cluster 1 showed a sharp connectivity spike under treatment that largely returned to baseline after recovery. We found enrichment for epithelial morphogenesis, cell migration, and cell adhesion GO terms, see Supplementary Figure S6A. Cluster 2 gained connectivity only in recovery and was enriched for ER and Golgi-associated pathways, including protein folding and transport (Supplementary Figure S6B). The third cluster maintained relatively stable, moderate connectivity across all conditions and resulted in a single enriched GO term (response to unfolded protein). Cluster 4 showed a sustained decrease in connectivity under treatment followed by a regain during the recovery phase. There, we detected enrichment for proteostasis-related processes, including ER stress and degradation pathways (Supplementary Figure S6D).

To illustrate these rewiring patterns, we selected 16 thyroid-relevant features representing all four clusters and reconstructed their condition-specific subnetworks in Cytoscape (Figure 6B). This visualization highlights pronounced differences in local connectivity and cluster composition across conditions. In the control network, the main connected component was largely composed of edges from clusters 2 and 4, reflecting coordinated baseline interactions within protein trafficking and degradation pathways. Under PTU treatment, features primarily assigned to cluster 1 expanded into a dense, highly connected subnetwork, whereas several control-state interactions were weakened or lost. During recovery, this treatment-associated component partially retracted toward intermediate connectivity. Cluster 4 edges re-emerged, indicating restoration of previously suppressed interactions, whereas cluster 2 edges contributed additional recovery-specific connections.

Together, these subnetworks illustrate a dynamic but partially reversible rewiring process, characterized by treatment-induced reorganization and recovery-associated restructuring rather than a simple return to the baseline state.

## Discussion

Integrative gene-correlation network analyses have become popular for combining multi-modal omics datasets, with a variety of strategies proposed for integrating transcriptomic, proteomic, and metabolomic layers [4, 5, 7, 9] Yet, preprocessing practices remain inconsistent and often rely on heuristic decisions. In this study, we reviewed existing methodologies and proposed practical guidelines for the preprocessing and integration of single-omics data. We demonstrated that standardizing each omics layer, whether individually or jointly, has no impact on the underlying correlation structure. Therefore, we advocate for a sample-wise concatenation of independently processed omics layers as the most efficient and robust approach for multi-omics WGCNA.

By applying this strategy, we constructed correlation networks for control, PTU-treated, and recovery conditions. The resulting modules contained features from all three molecular layers in proportions reflective of the input data, suggesting successful integration.

To examine how chemical exposure affects molecular network structure, we employed two complementary methods: (i) Module preservation analysis, to identify modules of co-regulated biomolecules that were present in one network but lost under treatment or recovery conditions, and *vice versa*. (ii) Differential network analysis, to identify features that significantly altered their connectivity across different networks.

Module preservation revealed that more than half of the control network modules were only weakly preserved under PTU treatment, indicating widespread disruption of baseline co-regulatory structures. Conversely, only two modules from the treated network showed weak evidence for preservation in the control network, suggesting that the adaptive structures present in the treated condition were predominantly subsets of pre-existing modules in normal conditions. This asymmetry indicates that PTU treatment primarily disrupts established regulatory patterns rather than inducing new, stable network structures, hence hinting at an adverse impact on the organism’s biological stability.

Differential connectivity analysis further highlighted the extent of disruption, as over 30% of all features (including all three biomolecule types) exhibited significantly altered connectivity in the control versus treated comparison. The majority of these differentially connected features (DCFs) lost connectivity under PTU treatment. Notably, a substantial subset of features showed evidence of re-wiring during the recovery phase, with over 3% demonstrating significant changes between treated and recovery networks. The absence of significantly altered connectivity between control and recovery network features suggests a general return to baseline connectivity. However, many features exhibited only partial restoration, potentially due to the limited recovery duration of two weeks following the four-week PTU exposure. This interpretation is reinforced by a strong negative correlation (Pearson’s *r* = −0.84) between *DiffK* values from the control-treated and treated-recovery comparisons, supporting the concept of reversible network disruption.

We implemented a feature-specific permutation-based p-value calculation for *DiffK*, enabling statistical inference on individual connectivity changes, a functionality not previously available in WGCNA-based differential network analysis. This extension increases confidence in identifying significantly rewired features and enables pathway-level assessments of differential connectivity, paving the way for more nuanced interpretations of regulatory network dynamics under external stressors.

In the integrative interpretation, we occasionally relate connectivity changes to previously reported shifts in molecular expression levels. However, differential expression analysis was not performed as part of the present study. Instead, expression results were taken from our prior work, in which the underlying transcriptomics, proteomics, and metabolomics datasets were generated and analyzed for differential expression [2].

### Differential Network Analysis Highlights Reversible Thyroid Perturbations

To gain deeper insights into specific pathways affected by PTU treatment, pathway analysis based on the *DiffK* metric identified several treatment-related alterations in the connectivity of molecular pathways and their features. Besides general stress response pathways such as ‘Cellular responses to stress’ and ‘Cellular responses to stimuli’ being significant, we took a detailed look at thyroid-specific perturbations—more specifically Na^+^/I^-^-Symporter (NIS) (de)regulation and pathway interactions.

NIS, which is encoded by *Slc5a5*, plays a fundamental role in thyroid hormone biosynthesis by mediating iodide uptake into follicular cells [52]. Its precise cellular localization and functionality rely on intricate transcriptional and post-translational regulatory mechanisms, ER-to-Golgi trafficking, glycosylation, membrane transport, and degradation pathways [53]. Differential network analysis revealed significant rewiring in these interconnected processes upon PTU treatment, providing a systems-level view of thyroid disruption and subsequent recovery.

A particularly relevant finding was the increased connectivity and upregulation of the ‘Signaling by NOTCH’ pathway during PTU exposure, where both central receptors, *Notch1* and *Notch3*, showed significant connectivity gains. Activated *Notch1* has previously been shown to upregulate *Slc5a5* directly by binding to its promoter [54], influencing broader thyrocyte-specific transcription programs [55]. Our analysis identified *Notch3* among the top 10 differentially connected features, further emphasizing its role in thyroid regulation. Besides its direct influence on transcription, Notch receptors and NIS share substantial similarities in their cellular processing: both are synthesized in the ER, undergo glycosylation within the Golgi apparatus, and depend on vesicle-mediated transport for membrane localization. Our analysis identified disruptions and subsequent network rewiring in pathways critical for these trafficking processes.

Elevated Notch receptors and NIS levels inherently increase the demand on glycosylation systems. Interestingly, the ‘Asparagine N-linked glycosylation’, covering the most important form of post-translational modification for proteins in the ER [56], showed a significant loss of connectivity during treatment followed by a restoration during recovery. For NIS, changes in the glycosylation pattern have been linked to decreased iodide transport activity [57]. Similarly, Notch signaling is severely affected by glycosylation disruption [58]. We also observe significant enrichment of three glycosylation pathways within the control lightcyan module, which exhibited only limited preservation in the network of treated samples, hence further indicating systemic disruptions of glycosylation processes.

Similar patterns were observed for trafficking-related pathways. Specifically, pathways like ‘ER to Golgi Anterograde Transport’, ‘Golgi-to-ER retrograde transport’, or ‘Vesicle-mediated transport’ significantly decreased connectivity under PTU treatment. The latter one on both transcriptome and metabolome layers. Notably, COPI-mediated transport, a critical component of NIS trafficking to the plasma membrane, showed a comparable trajectory of connectivity loss and recovery-facilitated restoration. *Arcn1*, encoding a COPI subunit, was among the top 10 significantly differentially connected features. Additionally, our results highlighted the roles of *Arf4* and *Vcp*, which experienced significantly decreased connectivity despite stable gene expression. *Arf4*, a small GTPase, is integral for NIS transport from the Golgi to the plasma membrane, while *Vcp* regulates NIS turnover, influencing whether NIS reaches the membrane or undergoes ER-associated degradation (ERAD) [59]. Reduced coordination between *Arf4* and *Vcp* could therefore critically impair NIS availability.

Endocytic processes further integrate the regulation of Notch receptors and NIS. Both proteins utilize canonical, clathrin-mediated endocytosis, impacting their cellular localization and functional availability. For Notch receptors, endocytosis regulates its activation, cleavage, and nuclear signaling, with unsuccessful membrane insertion limiting signaling efficiency [60]. Similarly, endocytosed NIS either recycles back to the membrane or is targeted for degradation, influencing its functional capacity [61]. Our analysis revealed a significant connectivity change in the `Reactome` pathway ‘Cargo recognition for clathrin-mediated endocytosis’ in the metabolomics layer. Furthermore, the gene encoding the clathrin heavy chain, *Cltc*, was also found to significantly lose connectivity with unchanged gene expression.

Taken together, these findings illustrate how differential network analysis can uncover complex yet coordinated molecular responses in transcriptional programs (Notch signaling), ER-Golgi dynamics, glycosylation, and endocytosis to regulate NIS functioning and localization. Interestingly, we observed several critical features, such as *Vcp, Arf4*, and *Cltc*, exhibiting stable gene expression but significantly reduced connectivity. Although counterintuitive at first glance, this pattern could indicate a selective, more focused cellular response under PTU-induced stress: these key proteins and their respective pathways might shift their functional interactions toward urgently needed survival pathways, effectively deprioritizing their broader cellular roles. Such a functional streamlining could ensure that essential pathways remain operational despite overall system-wide disruption. The divergence between stable expression and altered connectivity underscores that network reorganization can occur independently of differential expression, highlighting the complementary insight provided by differential network analysis.

### Global versus Local Alterations

The comparison of network alterations across experimental conditions using module preservation and differential connectivity (*DiffK*) analysis offers complementary perspectives on systemic perturbations. Module preservation assesses whether predefined modules of co-regulated features remain intact, whereas *DiffK* identifies individual features with altered co-expression regardless of their module affiliation. Below, we illustrate how the two approaches together reveal complementary regulatory changes across three biologically relevant domains impacted by PTU treatment.

#### Extracellular Matrix (ECM) remodeling

Pronounced changes in extracellular matrix (ECM) dynamics were observed under PTU treatment. *DiffK* analysis revealed significant enrichment of ECM-related pathways, including ‘Collagen degradation’ and ‘Assembly of collagen fibrils and other multimeric structures’ (`Reactome`), and ‘ECM-receptor interaction’ (`KEGG`), each exhibiting increased connectivity. Key ECM components, such as integrin *β*-1 (*Itgb1*) and collagen type IV (*Col4a1, Col4a2*), were notably among top connectivity gainers, suggesting treatment-induced stabilization or consolidation of ECM structures.

In contrast, ORA of the greenyellow control module, being the least preserved under PTU exposure, highlighted enrichment for glycosaminoglycan (GAG) degradation pathways, specifically ‘Glycosaminoglycan degradation’ (`KEGG`) and ‘Chondroitin sulfate/Dermatan sulfate degradation’ (`Reactome`). GAGs are essential structural components of the ECM, contributing to hydration, elasticity, and growth factor binding [62]. However, most GAG-related features showed connectivity loss, and only a few overlapped with differentially connected features from *DiffK* analysis. This suggests a disruption of GAG-associated remodeling processes that were not accompanied by a coordinated rewiring, potentially reflecting a decoupling from broader ECM adaptation.

Together, these results reveal a layered reorganization of ECM biology, wherein structural collagen networks gain coherence while GAG turnover appears suppressed or dysregulated. In general, thyroid hormones influence ECM composition, including fibronectin and collagens, across various tissues [63–65]. Previous studies have, furthermore, reported shifts in ECM remodeling across thyroid diseases, including Graves’ disease and thyroid cancer [66, 67], and GAG accumulation has long been associated with hypothyroidism-linked tissue swelling [68, 69].

#### Mitochondrial function

ORA identified pathways related to the tricarboxylic acid (TCA) cycle and oxidative phosphorylation in both the control (cyan module) and treated (turquoise) networks. Both modules were only moderately preserved when compared against the other condition, consistent with thyroid hormone roles in mitochondrial regulation [70, 71].

*DiffK* analysis did not identify mitochondrial or energy-related pathways as significantly enriched in the control versus treated comparison. Nevertheless, multiple core features (e.g., the genes *Ndufa7, Cox18*, and the protein LDHA) significantly lost connectivity under treatment, followed by a clear restoration during recovery. Additionally, the pathway ‘Citric acid (TCA) cycle and respiratory electron transport’ was significantly enriched on the proteome level in the treatment versus recovery comparison.

#### Lipid metabolism

*DiffK* highlighted substantial connectivity disruption in individual lipid metabolites. Particularly, 65 out of 68 significantly altered metabolites were lipids, including triglycerides, diglycerides, sphingomyelins, and hexosylceramides. Interestingly, all metabolites lost connectivity indicating strong network-level lipid disruption aligning with known PTU-induced disruptions of lipid homeostasis [72]. However, on pathway-level no lipid- or fatty acid-related pathways were enriched, likely due to limited coverage of structurally diverse lipid species in standard pathway databases [73].

Conversely, ORA identified enriched lipid metabolism pathways in modules from all three networks. The cyan (control) and darkturquoise (recovery) modules were dominated by triglycerides (63 and 46 features, respectively) and only moderately preserved under PTU treatment. In the treated network, triglycerides were more dispersed, particularly accumulating in the pink module, which alone contained nearly 200 lipid features and showed enrichment for diverse lipid-related pathways. Despite preserved lipid-centric modules, distinct reorganizations under stress suggest specialized lipid metabolism adaptations during chemical stress.

Taken together, these examples underscore how module preservation and differential connectivity analyses offer synergistic insights into network perturbations. Module preservation captures collective shifts in predefined co-expression patterns without directional inference, whereas *DiffK* provides higher resolution into feature-specific rewiring. While *DiffK* adds interpretive depth by identifying precise changes in local connectivity, it requires careful biological interpretation, as increased connectivity can reflect regulatory activation, network stabilization, or compensatory adaptation. Employing both strategies concurrently enhances both the sensitivity and biological interpretability of network-based assessments of systemic perturbation, hence anchoring global perturbations to molecular contexts.

### Local Network Reorganization and Adaptive Remodeling

Adjacency profiles around key thyroid-specific features, such as membrane transporters, membrane-associated signaling molecules, and trafficking or proteostasis regulators, offered a complementary, feature-centered perspective on network rewiring, building on the global and modular insights revealed by *DiffK* and module preservation analysis. Among the four clusters derived from clustering connectivity patterns across Control, Treated, and Recovery conditions, cluster 1 was enriched for epithelial morphogenesis pathways and exhibited a transient increase in connectivity during treatment. This enrichment is consistent with the known morphological changes of thyroid follicular epithelial cells under the influence of PTU [74]. Clusters 2 and 4 were enriched for ER- and Golgi-associated pathways, including protein folding, proteasome activity, and degradation processes but also trafficking processes. Notably, these changes were most prominent during the recovery phase, suggesting molecular effects that extend beyond the treatment period and may reflect adaptive remodeling rather than adverse disruption. These recovery-specific signatures were largely absent from our fold-change-based enrichment results [2], underscoring the ability of network-based approaches to uncover latent regulatory dynamics.

The derived network visualizations highlighted shifts in local connectivity and modular structure around thyroid-relevant features under treatment and recovery. In the control state, the thyroid molecular network exhibits clear compartmentalization, with minimal cross-talk between distinct membrane/ECM-associated proteins (*Slc5a5, Atp1b1, Col4a1*) and ER-trafficking components (*Arcn1, Vcp, Uso1, Cltc*). Following PTU exposure, the network underwent substantial reorganization, characterized by extensive cross-talk among previously isolated signaling, membrane transport, and ECM pathways—particularly integrating key regulators like *Notch3* and ECM components into a densely interconnected module. Interestingly, while membrane-associated, signaling, and ECM functions formed a highly mixed, adaptive module, the ER-trafficking and glycosylation machinery (*Arcn1, Arf4, Uso1, Vcp, Alg2, Cltc, Edem3*) remained distinctly compartmentalized and functionally coherent, albeit internally rewired. Upon recovery, partial segregation returned, indicating gradual reversion toward baseline compartmentalization. However, persistent rewiring, including renewed isolation of signaling nodes like *Notch3* and *Vwa1*, suggests the establishment of a shifted equilibrium rather than a full return to baseline within the 2-week recovery period.

### Multi-omics Integration Facilitated Biological Interpretation

Distinct omics layers capture different aspects of cellular regulation: transcriptomics reflects gene expression, proteomics reports on translational and post-translational control, and metabolomics represents downstream biochemical activity [1]. Consequently, specific biological signals may emerge uniquely in one layer or only become interpretable through integration.

In our analysis, lipid metabolism served as a prime example of this complementarity. Although pathway-level enrichment for lipid metabolism was largely absent, *DiffK* identified substantial rewiring among lipid metabolites at the feature level. This aligns with well-documented PTU-induced disruptions in lipid metabolism [72], which might have remained undetected using transcriptomics alone. Likewise, collagen and sphingolipid-related processes became evident only through a combined interpretation of module preservation and differential connectivity, despite their marginal appearance in individual omics analyses.

More broadly, multi-omics integration enhanced detection sensitivity by amplifying weak but consistent signals across layers. Using `multiGSEA` [23], which combines pathway-level p-values across omics types, we identified several biologically relevant pathways that did not reach significance in single-layer GSEA. For instance, ‘Collagen degradation’ and ‘PTEN Regulation’ became significant only in the integrated analysis, revealing cumulative regulatory shifts.

Despite differences in coverage, each omics layer contributed uniquely to the network analysis. Transcriptomics typically offers the broadest feature representation, while proteomics and metabolomics often cover fewer features, especially in legacy datasets or targeted techniques. As discussed in [2], limited coverage can restrict pathway detection and network resolution. Nevertheless, even smaller layers can yield disproportionately valuable insights, particularly when they capture regulatory processes that are inaccessible at other levels. Rather than being overshadowed by larger layers, their integration enhances the overall signal-to-noise ratio by leveraging cross-layer concordance and complementary biological information.

Together, these findings demonstrate that multi-omics integration offers a more holistic and sensitive view of molecular responses than single-layer analyses. In complex biological settings such as PTU-induced hypothyroidism, it enables the detection of nuanced regulatory changes that span molecular hierarchies, offering valuable insights for toxicological evaluation and mechanistic understanding.

## Conclusion

This study establishes a robust and interpretable framework for multi-omics co-expression network analysis using WGCNA. We propose practical guidelines for preprocessing and simultaneous integration of different omics layers. By integrating transcriptomics, proteomics, and metabolomics data into a unified network model, we improved detection of cross-omic regulatory interactions and highlighted the added value of multi-layer data integration in toxicological research. We introduced a statistically sound, permutation-based method to infer significance levels for differential connectivity at the feature level, providing reliable and interpretable insights into molecular rewiring. Combined with module preservation statistics, this dual strategy revealed condition-specific shifts both at the modular and feature scale, enabling comprehensive analysis of systemic perturbations. Our results demonstrate that applying such best practices enhances both biological relevance and interpretability, paving the way for more reproducible and mechanistic systems-level studies in the multi-omics era.

## Supporting information

**S1 Text. Correlation invariance under linear transformations**. Mathematical derivations showing the invariance properties of Pearson correlation and biweight midcorrelation under linear transformations.

**S1 Fig. Hierarchical clustering dendrogram of control, treated, and recovery samples**. Samples are labeled according to experimental group and duration. Panels show control (A), treated (B), and recovery (C) samples. Abbreviations indicate treatment duration, dose, and recovery phase.

**S2 Fig. Dendrograms for different groups**. Hierarchical clustering dendrograms for control (A), treated (B), and recovery (C) groups.

**S3 Fig. Association analysis of gene co-expression network modules with experimental, clinical, and histopathological traits**. Heatmaps show correlations between module eigengenes and external traits for control (A), treated (B), and recovery (C) networks. Asterisks indicate statistical significance.

**S4 Fig. Cross-tabulation heatmaps showing gene overlap between network modules**. Heatmaps compare module overlap across conditions: control versus treated (A), recovery versus treated (B), and control versus recovery (C). Cell values indicate overlapping genes, and colors represent −log_10_(*p*) values from Fisher’s exact test.

**S5 Fig. Module preservation analysis**. Median rank preservation statistics are shown for module sizes across six pairwise comparisons. Each point represents a module, and lower median rank indicates stronger preservation.

**S6 Fig. GO enrichment results for connectivity clusters (1–4)**. Enrichment results were obtained using over-representation analysis (ORA) with the `enrichGO` function. Panels (A–D) correspond to clusters 1–4, respectively.

**S1 Table. Significantly differentially connected pathways between control and treated conditions**. Reactome and KEGG pathways are listed with adjusted p-values for transcriptomics, proteomics, metabolomics, and the combined analysis.

**S2 Table. Significantly differentially connected pathways between treated and recovery conditions**. Reactome and KEGG pathways are listed with adjusted p-values for transcriptomics, proteomics, metabolomics, and the combined analysis.

## Footnotes

1 CEFIC-LRI C5-XomeTox project

2 https://www.ncbi.nlm.nih.gov/bioproject/PRJNA695243/

3 https://www.ebi.ac.uk/pride/archive/projects/PXD053208

4 http://dx.doi.org/10.21228/M8MD8N

5 https://edo98811.github.io/WGCNA_official_documentation/faq.html

6 DOI: 10.5281/zenodo.19592429

7 https://edo98811.github.io/WGCNA_official_documentation/faq.html

8 ORA_results_multiomics_WGCNA.zip

